# Reciprocity and behavioral heterogeneity govern the stability of social networks

**DOI:** 10.1101/694166

**Authors:** Roslyn Dakin, T. Brandt Ryder

**Author notes:** Corresponding authors: Roslyn Dakin. Corresponding authors: Brandt Ryder.

## Abstract

The dynamics of social networks can determine the transmission of information, the spread of diseases, and the evolution of behavior. Despite this broad importance, a general framework for predicting social network stability has not been proposed. Here, we present longitudinal data on the social dynamics of a cooperative bird species, the wire-tailed manakin, to evaluate the potential causes of temporal network stability. We find that when partners interact less frequently, and when the breadth of social connectedness within the network increases, the social network is subsequently less stable. Social connectivity was also negatively associated with the temporal persistence of coalition partnerships on an annual timescale. This negative association between connectivity and stability was surprising, especially given that individual manakins who were more connected also had more stable partnerships. This apparent paradox arises from a within-individual behavioral trade-off between partnership quantity and quality. Crucially, this trade-off is easily masked by behavioral variation among individuals. Using a simulation, we show that these results are explained by a simple model that combines among-individual behavioral heterogeneity and reciprocity within the network. As social networks become more connected, individuals face a trade-off between partnership quantity and maintenance. This model also demonstrates how among-individual behavioral heterogeneity, a ubiquitous feature of natural societies, can improve social stability. Together, these findings provide unifying principles that are expected to govern diverse social systems.

**Significance Statement:** In animal societies, social partnerships form a dynamic network that can change over time. Why are some social network structures more stable than others? We addressed this question by studying a cooperative bird species in which social behavior is important for fitness, similar to humans. We found that stable social networks are characterized by more frequent interactions, but sparser connectivity throughout the network. Using a simulation, we show how both results can be explained by a simple model of reciprocity. These findings indicate that social stability is governed by a trade-off whereby individuals can either maintain a few high-quality partners, or increase partner number. This fundamental trade-off may govern the dynamics and stability of many societies, including in humans.

---

Social network structure – or, the way individuals are linked by repeated social interactions – can influence the transmission of information, culture, resources, and diseases (1–6). Recent work has begun to demonstrate how changes to social network topology can have diverse costs (2, 7–10) and benefits (11–13), and may even influence the evolution of behavior (14). Although previous research has explored how social relationships form, and why some relationships are maintained for longer time periods than others (15–19), we lack a general framework to link individual behaviors with temporal dynamics of the social network at a collective level (20, 21). A mechanistic understanding of these processes is essential to explain the diversity of social systems, predict the downstream fate of social interactions, and engineer societies.

Here, we combine repeated measures of social structure and mechanistic models to elucidate the drivers of temporal dynamics in a cooperative system. Our empirical approach was based on automated biologging of a neotropical bird species, the wire-tailed manakin *Pipra filicauda*. Cooperative partnerships are a key part of the manakin social system (5, 22, 23), similar to humans (24). Male wire-tailed manakins cooperate by forming display coalitions which are the basis of dynamic social networks (25, 26). Cooperation occurs on display territories that are clustered into spatial aggregations called leks. A lek typically has between 4-14 display territories, and about twice as many males that visit the lek (each of whom limits his interactions to particular coalition partners (27)). The male-male coalitions that form at the manakin lek territories are a prerequisite for an individual to attain territory ownership, and ultimately, sire offspring (22, 23). To quantify cooperative partnerships and social network dynamics, we used an automated tracking system that identified times when two males co-occurred on a lek display territory as an indication of social interaction events (25, 26).

By tracking a population of 180 male manakins and 36,885 social interactions over three years, we took repeated measures of 11 leks to characterize the temporal dynamics of the social network at each site (on average, repeated measures of the same lek were 21 days apart; IQR 17-24). To analyze the dynamics of network topology from time *t*_1_ to *t*_2_, we define the stability of a binary network as the number of male-male partnerships (network edges) shared by both time points divided by the number of partnerships at either time point (i.e., the intersection divided by the union; Fig. 1a). To avoid bias as a result of rare events, we filtered the manakin data to include only significant partnerships in the computation of stability (see Methods for details). Using this metric of stability, we found that the manakin social networks were more stable than expected by chance (Fig. 1a), similar to other social animals (7, 28–30). However, stability was not constant, because each network fluctuated across a range of values (mean stability 0.43 ± SD 0.23; repeatability of stability 0, 95% CI 0–0.22).

**Fig. 1.**
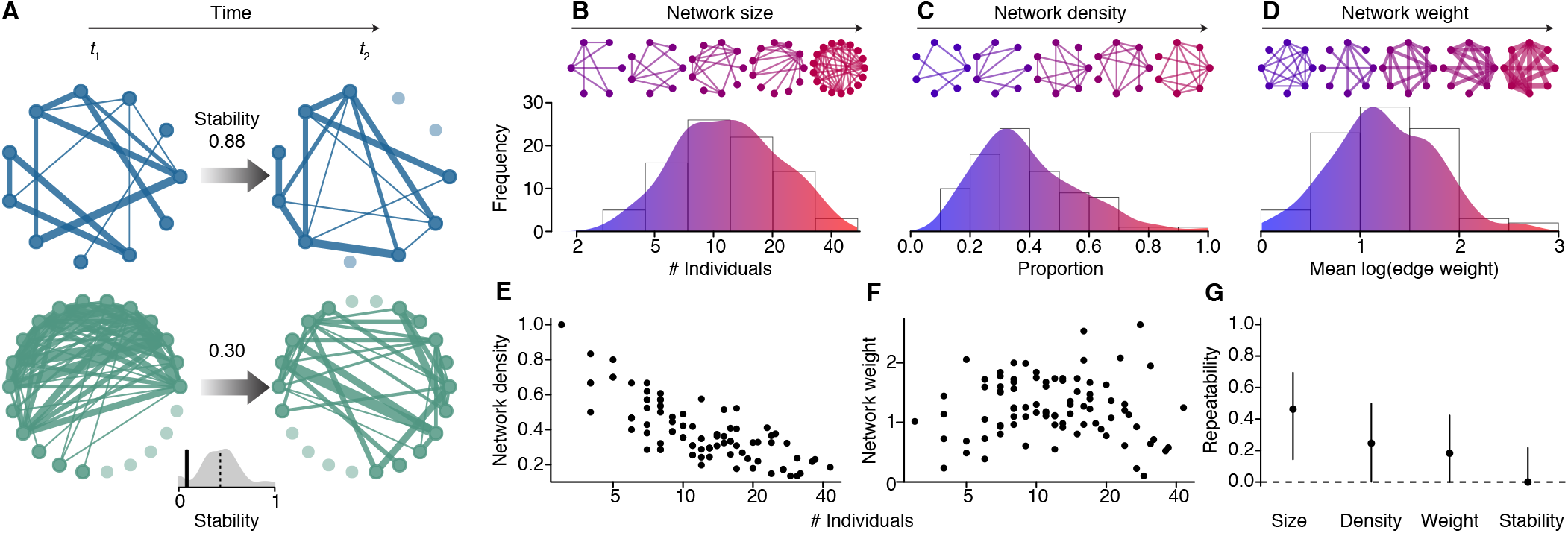
The temporal stability of social networks. The examples in (A) illustrate the definition of network stability. Two initial manakin social networks are shown (blue and green), with individuals depicted as nodes, and edge thickness weighted by the interaction frequency on a log scale. When the same two networks were sampled a second time, the edge structure of the blue network remained mostly stable, whereas the structure of the green network had largely changed. Manakin social networks were more stable than expected by chance, as shown by the fact that nearly all of the observed stabilities in the grey distribution exceed the 95% confidence interval of the null expectation (vertical black bar). The observed networks also varied in properties such as: (B) the number of individuals in the social network (size), (C) the proportion of possible relationships that occurred (density, a measure of connectivity), and (D) the average frequency of interactions (weight). In the illustration for network weight, edge thicknesses are also scaled to the average interaction frequency. (E-F) Scaling of density and weight with network size. (G) Repeatability of network properties ± 95% confidence intervals (n = 86 repeated measures, 60 for stability, of 11 lek networks).

To test how the social structure at *t*_1_ might predict subsequent network stability, we used a mixed-effects modelling framework. We found that three properties of the social network could explain 28% of the variation in stability in the best-fit model: network size (the number of individuals or nodes in the network), network weight (the average frequency of social interactions), and network density (the proportion of possible partnerships that actually occurred, which is a measure of connectivity; Fig. 1b-d). Note that network size, weight, and density were all determined using unfiltered weighted networks (see Methods for details). The analysis also accounted for the timing (year and mean Julian date) when each sample of a network was taken. All else being equal, when partners within the network interacted more often (higher network weight), and when there were relatively fewer partnerships in the network (lower network density), the social structure was more stable over the subsequent weeks (Fig. 2).

**Fig. 2.**
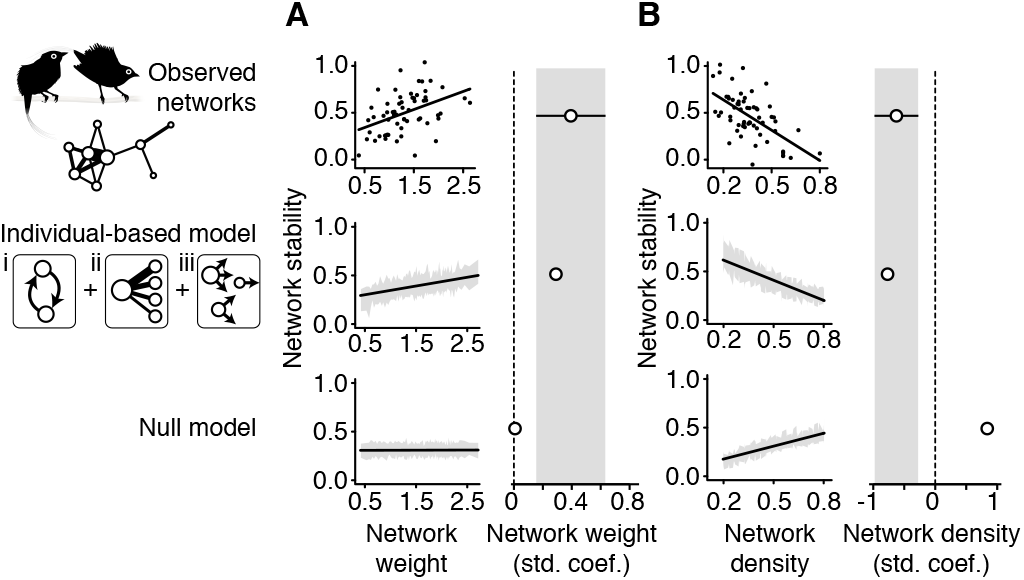
A model of reciprocity and behavioral heterogeneity predicts network stability. (A-B), The stability of a social network is positively associated with the average frequency of interactions (weight), whereas stability is negatively associated with the relative density of network connections. These effects were confirmed in an individual-based simulation of reciprocity that combined three behavioral rules: (i) a requirement for reciprocal partner choice, (ii) a preference for previous partners, and (iii) repeatable variation among individuals in social behavior. The left columns in A and B show partial residual scatterplots from the statistical analyses, after accounting for additional covariates (Tables S1-S3; n = 60 for the observed networks and n = 3,000 for the simulations). Because the simulation sample sizes are so large, shading is used on the simulation scatterplots to show the 95% central range of data binned along the x-axis instead of plotting individual points. The right columns in A and B show the standardized effect sizes (± 95% confidence intervals for the observed networks; these intervals are also extended with shading for direct comparison with the simulations). The coefficients derived from the individual-based model fall within the 95% confidence intervals of the observed data, unlike the null model (which was a simulation with rules i-iii removed). Note that the 95% confidence intervals for all simulation effect sizes are not shown because they are narrower than the data points.

The negative association between network density and stability was unexpected, given that connectedness in a cooperative system without defection is thought to foster social cohesion (31). To provide a mechanistic explanation for this result, we built a simulation model based on the hypothesis that individual behaviors would drive emergent properties of the system (32, 33). In this model, the individuals iteratively sought partnerships with each other at each time step. The model assumed three simple rules that describe a scenario of reciprocity (24) with among-individual heterogeneity (32, 34): (i) social partnerships are formed through reciprocal partner choice, wherein both individuals must choose each other; (ii) individuals prefer social partners with whom they have previously interacted (35); and (iii) there are consistent among-individual differences in the expression of social behaviour. This third assumption of among-individual behavioral heterogeneity is ubiquitous in human and animal behavior (34) and has been shown to influence collective performance (36) and the evolution of cooperation (37, 38). It is important to note that our simulation model made no assumptions as to the source of this among-individual heterogeneity (which could be caused by genetic, environmental, age-related, or other factors). We ran this simulation on 3,000 initial networks that were generated *de novo* to represent a broad range of network sizes, weights, and densities, and we used these initial networks to parameterize (ii) and (iii). We then allowed the individual nodes to repeatedly interact with each other. Finally, we computed the stability of each simulation run, by comparing the initial structure to the one that resulted from the newly simulated interactions.

Similar to the manakin data, we found that networks with a relatively high frequency of cooperation (high weight) but sparse connectivity (low density) were more stable (Fig. 2). Hence, the simple model of reciprocity plus heterogeneity was sufficient to recreate the dynamics observed in the empirical networks. Moreover, we found that the null model simulations that lacked all three assumptions (i-iii), or that included only (i), (ii), or (iii) alone, were insufficient to generate the empirical patterns of stability. In null models that lacked reciprocity, denser networks were also consistently more stable, making the negative effect observed in the empirical data particularly striking (Fig. 2b). Overall, our findings indicate that both behavioral processes, reciprocity and heterogeneity, are necessary to recreate the weight and density effects on network stability. Finally, we found that the larger simulated networks with more individuals were also significantly less stable, independent of network weight and density. This effect of network size was also consistent with the manakin data (although in the empirical analysis, it was not quite statistically significant; Table S3).

Why are some social partnerships able to persist through time (7, 18, 29, 30)? To understand how social structure might influence the fidelity of particular bonds over longer timescales, we analyzed the annual persistence of 669 manakin partnerships from one season to the next (Fig. 3a-b). In this analysis, a partnership was defined as two males who interacted on a display territory at least once in a given season. Annual persistence was defined as that partnership recurring at a significant rate the following year (see Methods for details). The analysis accounted for the identities of the partners, the year, the lek where the partnership occurred, and other factors including the spatial overlap of the individuals. Two features predominantly explained the variation in partnership persistence: the interaction frequency (edge weight), and the local social density (edge connectivity, which quantifies the number of alternative paths that can connect two partners in a social network). Specifically, a partnership was more likely to persist if the two individuals interacted more frequently, but had lower connectivity in their social neighborhood. These results are consistent with the phenomena observed at the network level over shorter weekly timescales (Fig. 2). Moreover, we found that the simulation of reciprocity and heterogeneity could also recreate the empirical results found for partnership persistence (Fig. 3c-d).

**Fig. 3.**
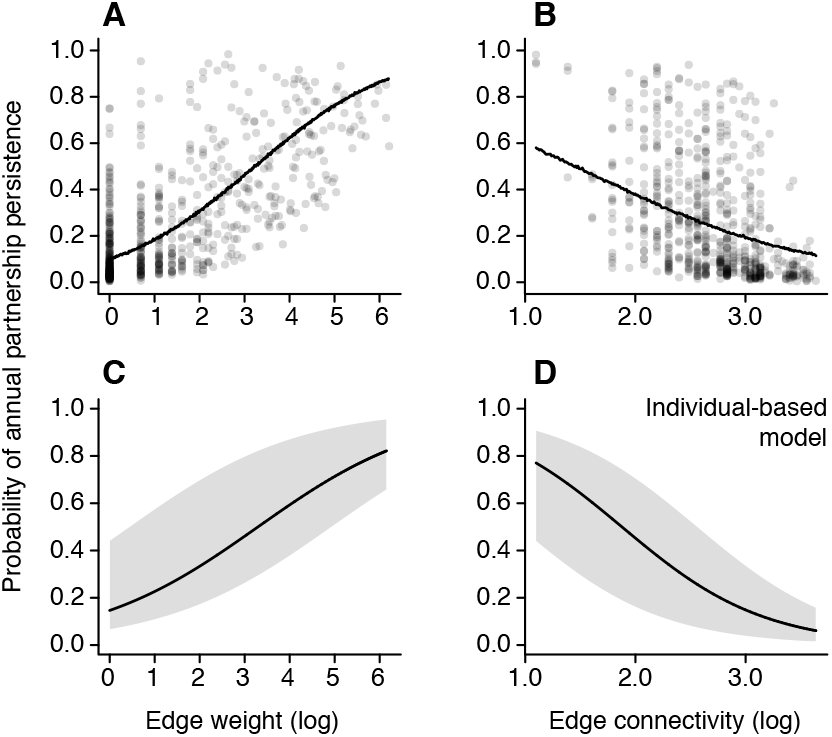
Social structure predicts the long-term persistence of social partnerships. (A-B) The probability that a partnership persisted across years was greater when the two partners interacted at a higher frequency (edge weight), but had fewer alternate paths connecting them in the social network (edge connectivity). Data points show how edge weight and connectivity (x-axes) determine the predicted probability of partnership persistence (y-axis) in a multiple-membership analysis (n = 669 partnerships among 91 individuals). (C-D) The influence of edge weight and connectivity is also found in the individual-based model of reciprocity described in Fig. 2. Shaded areas in C-D show the 95% central range for partial residuals binned along the x-axis.

These negative effects of overall network connectivity suggest that social stability is governed by a fundamental trade-off between the quantity and quality of social partnerships. Contrary to the trade-off hypothesis, however, the manakins with more partners (i.e., those with higher average degree centrality) formed coalitions that were more likely to persist through time (Fig. 4a). This apparent paradox is resolved by partitioning the variation among- and within-individuals (Fig. 4b-c). Among individuals, the males who were more connected were better able to maintain their partnerships (Fig. 4b). However, when a given male had more partners than his average, he was less able to maintain them (Fig. 4c). Thus, each individual may have a different threshold for the number of stable coalition partnerships he is able to maintain. This explains why densely connected social networks are less stable (Fig. 2), even though well-connected individuals are better at maintaining partnerships (Fig. 4a-b). In wire-tailed manakins, the proximate causes of this among-individual heterogeneity are not yet well understood (39), but could include a male’s quality, age and social experience, and/or his compatibility with the other males on his lek.

**Fig. 4.**
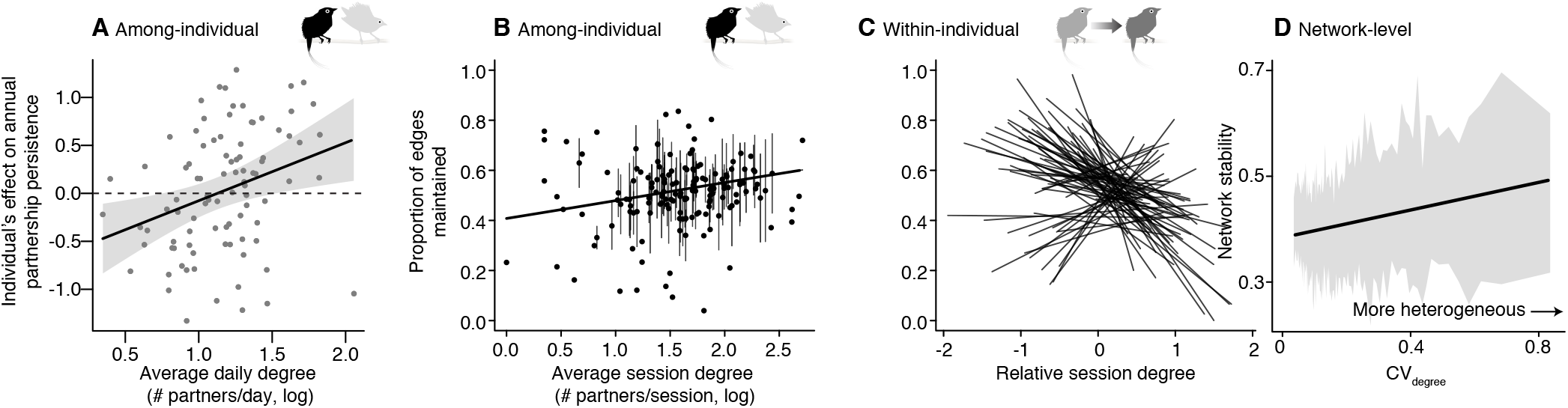
Behavioral heterogeneity and social stability. (A) The males who consistently interacted with more partners per day (high average daily degree, x-axis) promoted long-term coalition persistence (y-axis; n = 90 individuals; ± 95% confidence interval of the prediction line). (B-C) However, a trade-off is revealed when examining repeated measures within-individuals. The plots in B and C show an analysis of within-season partnership maintenance (n = 565 repeated measures of 152 individuals). Despite the positive among-individual effect shown in B, at times when a given male had more partners than his average in c, he was less able to maintain stable partnerships. To visualize among-and within-individual variation, a single average is plotted for each male in B (± SE if a male had >3 measurements), whereas a separate linear fit is shown for individuals with >3 measurements in C. (D) In a simulation model, social networks with greater among-individual behavioral heterogeneity (CV_degree_) were also more temporally stable. The y-axis shows partial residuals from an analysis that also accounts for the effects of network size, weight, and density (n = 3,000). Shading indicates the 95% central range for partial residuals binned along the x-axis.

How might the magnitude of behavioral heterogeneity influence the stability of cooperative networks (36)? Our simulation model provided an opportunity to begin exploring this question. To measure heterogeneity, we computed the coefficient of variation in degree centrality (CV_degree_) in each of the initial networks; higher values indicate greater behavioral heterogeneity in the system (40). We found that CV_degree_ had a significant positive effect on subsequent network stability (Fig. 4d), demonstrating that individual variation in sociality can foster stable social networks. This is similar to the way some ecological systems are affected by heterogeneity (e.g., CV of connectedness (degree) and edge weights) (40, 41). In social systems, behavioral heterogeneity can also include suites of correlated traits such as dispersal, risk-taking, and cognitive ability, in addition to variation in sociality (20, 34, 42). Further study is needed to understand how this covariation influences social network stability and the evolution of complex social behavior (33).

In summary, we find that social interactions can have opposing effects on the stability of cooperative systems. On the one hand, the stability of the social network is enhanced by increasing the interaction frequency among a small number of partnerships. However, when individuals become too broadly connected, the social network can be destabilized. This is because individuals are constrained in their ability to reciprocate a large number of social partnerships. Our results also highlight the fact that among-individual heterogeneity can easily mask this behavioral trade-off (34). Hence, these results emphasize the importance of longitudinal data that captures multilevel variation, within- and among-individuals.

Can these principles be applied to other systems? Although social network stability has not yet been analyzed in humans at a broad scale, this is an important next step, given that globalization and social media use have rapidly increased the breadth of human social connectivity (6, 43). Our model provides one potential explanation for how these novel behavioral interaction patterns could have a destabilizing effect on human social structure. Another important question is how much topological changes in these networks affect other dynamics, such as the spread of emotions, cultural evolution, and disease transmission. Although our study focused on one type of cooperative system, many other social networks are formed as a result of competitive, aggressive, mating, and information-sharing interactions (20). As a unifying framework, we propose that social stability in these other contexts will also be determined by the simple behavioral processes that generate heterogeneity, partner preferences, and the symmetry of partner choice.

## MATERIALS AND METHODS

### Field methods

Observed social networks were based on a study of wire-tailed manakins, *Pipra filicauda*, at the Tiputini Biodiversity Station in Ecuador (0° 38’ S, 76° 08’ W, 200 m elevation). Male wire-tailed manakins perform cooperative courtship displays at exploded leks, where males are in acoustic but not visual contact (44). The population at Tiputini has been monitored since 2002 to study the fitness benefits of cooperative behavior (22, 23). The present study spanned three field seasons (December-March) in 2015-16, 2016-17 and 2017-18, and used an automated proximity data-logging system to record cooperative interactions among males (25, 26). Manakins were captured using mist-nets and each male was outfitted with unique color bands and a coded nano-tag transmitter (NTQB-2, Lotek Wireless; 0.35 g). To record the social network at a given lek, proximity data-loggers (SRX-DL800, Lotek Wireless) were deployed in each territory to record all tag detections within the territory from 06:00 to 16:00 for ~6 consecutive days (± SD 1 day), which comprised a single recording session (26, 39). Territory ownership was assigned using direct observation of color-banded males at the display sites (22), and was subsequently verified in the proximity data. Sample sizes were not predetermined because our aim was to track all individuals within the studied leks (39). In the absence of a formal mark-recapture protocol, we examined the percentage of territory-holders tagged as an indication of how well our sample covered the known population (95%, 95%, and 92%, for the three respective field seasons). All animal research was approved by the Smithsonian ACUC (protocols #12-23, 14-25, and 17-11) and the Ecuadorean Ministry of the Environment (MAE-DNB-CM-2015-0008).

### Data processing

All data processing and statistical analyses were performed in R 3.5.1 (45). Male-male cooperative interactions on the display territories were determined using spatiotemporal overlap of tag detections in the proximity data (26). Specifically, a social interaction was defined as a joint detection of two males within approximately 5 m on a territory during the breeding season (26). This spatial range corresponds to the visual and acoustic contact required for a typical display interaction in this species (22, 46). Because the social interactions were measured using an automated system, the networks were constructed blind to the sociality of particular individuals and/or leks. A previous validation study conducted in 2012 (25) confirmed that the social interactions defined by this automated system corresponded to direct observations of male-male display partnerships. We used the social interaction data to build undirected weighted social networks for each lek recording session, with each node representing a male, and the edges weighted by the frequency of social interactions summed over a recording session (approx. 6 days, defined above). In total, we characterized 86 repeated measures of the social networks at 11 leks (mean 7.8 sessions per lek, ± SD 3.7) from 29,760 sampling session hours and 36,885 unique social interactions among 180 individuals. We used a clustering analyses in the igraph package (47, 48) to verify that our sampling design was well-matched to the inherent social structure of the population (Fig. S1).

### Network stability

The stability of social network topology is determined by both the gain and loss of associations over time. We therefore defined a bidirectional metric of social network stability for binary (unweighted) networks that compares two repeated measurements of the network, *N*, at times *t*_1_ and *t*_2_. The stability of *N* over the period *t*_l_ ↔ *t*_2_ is defined as the number of social partnerships (i.e., edges) shared by *N*_1_ and *N*_2_ (i.e., intersection ∩), divided by the total number of unique edge connections in either *N*_1_ or *N*_2_ (i.e., union ∪). Using *E* to represent network edges, stability is thus defined by the following formula:

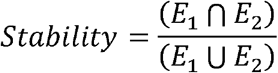

This metric can range from 0 (unstable) to 1 (highly stable). Note that this definition would not apply to complete (fully connected) networks. In most social networks, individuals (or nodes) can also be gained or lost over time, which alters the set of possible interactions that could occur. To ensure that our measure of social network stability was based on edges that could have occurred at both time points, only individuals who were present at both *t*_1_ and *t*_2_ were included in the calculation (49). Therefore, this definition captures the stability of relationships among individuals who remained in the network over two consecutive time steps (49). Furthermore, to ensure that the stability metric was not biased by rare interactions (50), we also filtered the stability calculation to be based on binary networks that included only edges that met two criteria in the empirical data: (1) a significant edge had to occur more often than its own average occurrence in 1,000 random permutations of the interaction data, and (2) it had to occur at least six times during the recording session (i.e., on average, about once per day). The second criterion ensured that rare interactions were not easily deemed significant. The value of six was chosen to correspond to the average number of days in each recording sessions, but we also verified that other thresholds >2 did not influence our results. Finally, we verified that all of the results were also unchanged when using only the second (absolute) criterion.

The average stability score for the manakin networks was 0.43 (± SD 0.23, n = 60 networks at 11 leks). Note that the sample size of 60 is smaller than the total number of recording sessions, because the stability dataset is limited to networks that were also sampled at *t*_2_ within the same season. The observed networks were also more stable than expected by chance (paired t-test, t = 12.08, p < 0.0001), as determined by random network rewiring (100 edge permutations for each of the 60 measurements; grand mean null expected stability 0.07 ± SD 0.05).

### Network-level analyses

Network size, connectivity, and structure can all influence the dynamics and stability of diverse network types (40, 41). Therefore, to determine how network-level properties at *t*_1_ predict subsequent social network stability, we fit mixed-effects regression models using the package lme4 (51) (n = 60 networks at 11 leks). The analysis included lek identity as a random effect, and to account for potential temporal trends, we also included field season (categorial) and mean Julian date of the network (continuous) as two fixed effects. Mean Julian date for a network was calculated as the average date of all of the social interactions that occurred within that network. We considered five network properties that have been shown to influence network dynamics in other contexts (40, 41) as additional fixed effects: (1) network weight is a measure of the average relationship frequency, calculated as the mean of the log-transformed edge weights; (2) network density is a measure of the breadth of connectivity, calculated as the proportion of relationships that actually occurred relative to a completely connected network; (3) clustering coefficient, or network transitivity, is an alternative measure of connectivity that is often important in social networks (52), and that describes the probability that a given individual/node’s social partners are also connected; and (4) network modularity is yet another measure of connectivity that describes how well the network can be subdivided into separate communities using the random-walk algorithm (47, 48). To account for the fact that these network-level properties often scale with network size (32, 53) (Fig. 1e-g), we also included (5) the log-transformed number of individuals/nodes in the social network as an additional predictor. Note that unlike the calculation of network stability in the previous section, all five of these network properties were computed from unfiltered network data.

Because density, transitivity, and modularity were all similar measures of network connectivity, and because the sample size of 60 is not enough to reasonably estimate more than five or six fixed effects at a time, we used a model selection procedure to compare candidate models that included field season, date, network size, and at most two of the other network properties. Given that network weight, density, and clustering coefficient were all correlated measures of connectivity, each model could include at most one of the those parameters. We also considered a null model that included none of (1)-(4). Complete details are provided in Tables S1-S2. Finally, we evaluated whether network stability was influenced by two logistical factors: first, sampling effort, and second, a testosterone manipulation experiment that was conducted for a separate study in 2016-17 and 2017-18 (n = 9 individuals out of 180 that were implanted with testosterone (39)). To verify that these two logistical factors did not influence our results, we added additional fixed effects for the number of recording hours (median 75, mean 73 ± SD 10) and/or the number of hormone-manipulated individuals in a given network (median 0, mean 0.10 ± SD 0.41), neither of which had a significant effect on network stability (all p > 0.43). We also verified that all of the conclusions of the network-level analysis were unchanged when accounting for either or both of these covariates. To determine the repeatability of network properties of the leks, we calculated the proportion of total variation that was due to differences among the leks using mixed-effects models with lek as the random effect and field season and Julian date as fixed effects (51, 54).

### Edge-level analysis

The edge-level analysis examined the persistence of manakin social partnerships on an annual timescale. This analysis considered 669 dyadic partnerships among 91 individuals wherein both individuals in the partnership were also present and tagged in the subsequent breeding season. A partnership was defined as two males who had interacted on a display territory at least once. The binary response variable, partnership persistence, was defined as whether a partnership was sustained and significant in the subsequent breeding season (using the criteria for significance defined above in the section “Network stability”). Because both individuals in a social partnership can contribute to its fate, and because they both had other partnerships in the dataset, we modelled persistence using a multiple-membership structure in a binomial mixed-effects regression model, fit with the brms package (55). This method can be used to account for multiple partner identities within a single random effect (26, 55–57). In our analysis, the two identities were weighted equally, because we assumed they could both determine partnership persistence. An additional random effect was included to account for the lek where each partnership occurred. The analysis also included fixed effects to account for the initial field season (categorical), the territorial status of the pair (categorical; either two territory-holders, a territory holder plus a floater, or two floaters (22)), the sampling effort at that lek in both the initial and the subsequent field season, and the initial spatial overlap of the pair, which can influence the probability of interaction (29). Because manakins use discrete display territories, we defined the spatial overlap of two males as the log-inverse of the chi-squared statistic comparing their distributions of territory detections (pings) in the proximity data; larger values of this metric indicate greater spatial overlap.

Based on the results of the network-level analysis, we sought to test whether edge-level network properties would predict partnership persistence. Thus, we also included the following fixed effects: (1) edge weight, or the log-transformed social interaction frequency; (2) edge betweenness, a measure of social centrality, defined as the log-transformed number of shortest paths passing through that edge; and (3) edge connectivity, a measure of social density, defined as the minimum number of edges that must be removed to eliminate all paths between the two individuals/nodes in a partnership (48). Edge weight and edge connectivity also correspond to the metrics of network weight and density, respectively, at the network level. In contrast, edge betweenness captures a different property: a relationship with a high edge betweenness is one that links individuals from two disparate communities. If partnership maintenance is enhanced when both individuals have strong links to the same local community, we expect a negative relationship between edge betweenness and persistence. Alternatively, if individuals place particular value on long-range ties, partnership persistence might be positively related to betweenness. We ran four independently seeded chains with default priors, storing 2,000 samples from each chain, and verifying that the convergence statistics were all equal to one (55) (Table S4).

### Among-individual analysis

To test whether partnership persistence could be attributed to behavioral differences among individual manakins, we refit the analysis described above, but without accounting for (1)-(3) listed above. The random intercepts from this model provide an estimate of among-individual variation in social stability (26, 54). We hypothesized that the following behavioral phenotypes (26) could affect this trait: (1) a male’s average daily effort, measured using his log-transformed count of detections (pings) on the leks; (2) his average daily strength, using his log-transformed sum of interaction frequencies; (3) his average daily degree, using his log-transformed number of social partnerships, and (4) and his average daily social importance, defined as the exclusivity of his partnerships (see the previous protocol (26) for additional details). Because these four phenotypes were also correlated (26, 39), we compared six candidate regression models, four of which included only one behavioral phenotype, one of which included all four phenotypes, and one of which included no behavioral phenotypes (n = 91 individuals; see Table S5). All candidate models included a male’s status as either a territory-holder or floater.

### Quantity-quality trade-off analysis

We next sought to test the hypothesis that individuals in a network face a trade-off between the quantity (number of partners) and stability of their social partnerships. Because among-individual variance can mask trade-offs that occur within-individuals (58), testing this hypothesis requires a variance-partitioning approach. To achieve this, we defined repeated measures of individual partnership maintenance as the proportion of a male’s coalition partners that were maintained from a given recording session to the next recording session (n = 565 repeated measures of 152 individuals). Similar to our other analyses, a partnership was defined as two males having at least one interaction during a recording session. Note that a male had to be present, tagged, detected, and not part of the hormone manipulation experiment in both the initial and subsequent recording sessions to be included in this sample. We used within-group centering to partition the variation in the predictor variable, degree centrality, within- and among-individuals (59). The first step was to determine log-transformed degree for each male in each recording session; next, we took a single average degree value per male; and finally, we calculated relative degree in each recording session as a male’s log-transformed degree minus his overall average. Thus, average and relative degree represent two orthogonal predictors that can be analyzed within the same regression model. The analysis was fit as a binomial mixed-effects model in lme4 (51) with a random effect of individual identity, and it also included two categorical fixed effects to account for field season and territorial status, respectively, as well as a continuous fixed effect to account for sampling effort (Table S6).

To evaluate what would be expected in this analysis by chance alone, we repeated the analysis using randomly permuted networks. To do this, we used permutations of the manakin data wherein each lek social network was randomly rewired between each recording session (48). We generated 1,000 of these randomized datasets and then performed the same repeated-measures analysis that was applied to the observed data. We averaged the results across all 1,000 randomized analyses to derive the null expectation shown in Fig. S2.

### Individual-based simulation models

To provide a mechanistic explanation for how individual behavior scales up to influence social network stability, we developed a simple individual-based simulation model. The model was based on the general principles of social reciprocity (24) and among-individual behavioral heterogeneity. There were three core assumptions: (i) individuals had to actively choose each other in order to form a partnership; (ii) each individual had a ranked set of preferences for social partners, predicted only by its previous social interaction frequencies in the initial network, and (iii) individuals expressed consistent differences in their social behavior (referred to as behavioral phenotype). The second rule (ii) is supported by strong evidence that social relationships are non-random and persist over long time-scales in human and nonhuman animals (7, 29). Together, rules (i) and (ii) also represent a form of reciprocal altruism (24), because prior interactions increase the probability that a partner will be re-chosen. Rule (iii) represents a phenomenon that is often referred to as among-individual variation, heterogeneity, or personality; it has empirical support across vertebrates (34), including in manakins (26).

To experimentally test the effects of network size, weight, and density on network stability, we generated 3,000 initial networks with diverse properties that were within the range of the observed data. Network size was first chosen from the range of 11-20 individuals or nodes (10 size bins). To manipulate network density along the same range observed in the manakin data, we first generated completely connected networks, and then randomly removed edges until a target initial density was achieved (targets ranging from 0.2-0.8, for a total of 20 target density bins). To generate a broad range of initial network weights, each edge weight was first sampled from the manakin data, and then multiplied by a weight constant ranging from 0.2-2.0 (15 weight factor bins). The resulting edge weights were then rounded up, to a maximum of 500. We generated 3,000 networks with all possible combinations of these network properties (10 × 15 × 20 = 3,000).

The simulation proceeded as follows. First, to satisfy rule (ii), we assigned a set of preferences to each node based on that node’s partnerships in the initial starting network. The set of preferences included all other nodes, ranked by interaction frequency with the focal node in the initial network. Hence, the probability of choice was correlated with initial interaction frequency. To satisfy rule (iii), each node was also allotted a specific number of interaction attempts per time step (ranging from 1-4). This number was calculated by log-transforming the strength of the focal node in the initial network (also referred to as weighted degree) to obtain its behavioral phenotype; higher values meant that a node could attempt more social interactions per unit time. To satisfy rule (i), a partnership was only formed if both nodes chose each other within a given time step. The simulation ran over five time steps and the final network was determined by summing the new interactions that occurred (Fig. S3). No filtering was applied to calculate network stability in the simulation. Note that for simplicity, the preference ranks for (ii) were not updated during the time steps that occurred within the simulation.

For the null model, we followed the same procedures above, except that each individual’s partner choice probabilities were assigned randomly to the set of all other nodes, the number of attempted interactions per time step was fixed across individuals, and reciprocal partner choice was not required for partnership formation in the null model (i.e., assumptions ii, iii, and i were removed). We also tested models with either (i), (ii), or (iii) alone. After running the simulations, we used linear models to statistically analyze the variation in network stability and examine the three predictors of interest from Table S2: network size, weight, and density. To compare the results of this analysis with the statistical estimates derived from the observed data, all predictors and response variables were standardized to have a mean of 0 and SD of 1 (Table S3). To test whether the simulation model of reciprocity and heterogeneity could also explain our edge-level analysis, we used a binomial mixed-effects regression of edge persistence in the simulation, with the identity of the initial network as a random effect, and edge weight and edge connectivity as the predictors.

We chose five as the number of time steps in these simulations to correspond to a period of about five days of behavioral activity. To verify that the results of the simulation model would be robust to alternative time parameters, we also repeated these analyses using simulations with either three or ten time steps instead. In each case, we reached the same conclusions with nearly identical effect sizes for network size, weight, and density, respectively (Table S3).

## Data deposition

The data and R code necessary to reproduce our results are available at: https://figshare.com/s/470aeac186a9dab72860

## Acknowledgements

We thank Ben Vernasco, Camilo Alfonso, Brent Horton, Ignacio Moore, Brian Evans, David and Consuelo Romo, Kelly Swing, Diego Mosquera, Gabriela Vinueza, the Tiputini Biodiversity Station of the Universidad San Francisco de Quito, Julie Morand-Ferron, Jean-Guy Godin, and three anonymous reviewers. Funding was provided by National Science Foundation (NSF) IOS 1353085 and the Smithsonian Migratory Bird Center.

