## Supplementary material for "Reciprocity and behavioral heterogeneity govern the stability of social networks": SI Appendix

**Table S1. Model selection for predictors of social network stability.** Candidate models to predict the stability of manakin social networks (n = 60 networks at 11 leks). Linear mixed-effects regression models were ranked by AIC. All models included a random effect of lek identity to account for repeated measures of the same lek, as well as fixed effects to account for annual/season variation (categorical with three levels: 2015-16, 2016-17, 2017-18), mean Julian date (continuous), and the size of the network (continuous; log-transformed number of individuals). The evidence ratio compares the support for model i to the best-fit model, and is calculated using the Akaike weights as $w_1/w_i$. $R^2_{LMM(m)}$ provides an estimate of the proportion of total variance explained by the fixed effects, whereas $R^2_{LMM(c)}$ is the variance explained by the fixed and random effects together (Nakagawa and Schielzeth 2013).

| Rank, i | Fixed effect predictors<br>Field season + Julian date + Network size…. | $AIC_i$ | $\Delta AIC_i$ | $w_i$ (AIC) | Evidence ratio<br>(best-fit vs. model i) | $R^2_{LMM(m)}$ | $R^2_{LMM(c)}$ |
|---|---|---|---|---|---|---|---|
| 1 | … + Network weight + Network density | 23.96 | 0 | 0.71 | – | 0.28 | 0.28 |
| 2 | … + Network weight + Clustering coefficient | 27.01 | 3.05 | 0.15 | 4.60 | 0.25 | 0.25 |
| 3 | … + Network density | 27.78 | 3.82 | 0.10 | 6.77 | 0.16 | 0.16 |
| 4 | … + (null) | 32.46 | 8.50 | 0.01 | 70.15 | 0.05 | 0.05 |
| 5 | … + Network weight | 32.50 | 8.54 | 0.01 | 71.50 | 0.14 | 0.14 |
| 6 | … + Clustering coefficient | 32.85 | 8.89 | 0.01 | 85.12 | 0.09 | 0.09 |
| 7 | … + Network weight + Modularity | 34.10 | 10.14 | 0.00 | 159.13 | 0.16 | 0.16 |
| 8 | … + Modularity | 34.82 | 10.86 | 0.00 | 228.34 | 0.07 | 0.07 |

**Table S2. Best-fit model of social network stability.**

Fixed-effect estimates from a linear mixed-effects model of network stability with a random effect of lek (n = 60 observed networks at 11 leks). The response variable, stability, and all continuous predictors were mean-centered and standardized (mean = 0, SD = 1) so that the estimates would be comparable within this analysis, and with the analyses of other simulated datasets.

| Fixed effect | Estimate (std. coef.) | Lower 95% confidence interval | Upper 95% confidence interval | t | p-value |
|---|---|---|---|---|---|
| Field season | | | | | |
| 2016-17 | 0.15 | –0.48 | 0.78 | 0.47 | 0.64 |
| 2017-18 | –0.18 | –0.77 | 0.40 | –0.61 | 0.54 |
| Julian date | 0.00 | –0.24 | 0.24 | 0.00 | >0.99 |
| Network size | –0.29 | –0.65 | 0.08 | –1.53 | 0.13 |
| Network weight | 0.39 | 0.15 | 0.63 | 3.22 | 0.002 |
| Network density | –0.63 | –0.98 | –0.27 | –3.47 | 0.001 |

**Table S3. Analyses of network stability in individual-based simulation models.** We examined the network-level properties that predict stability when partnerships are governed by individual (node) behavior. Six linear regression analyses are shown here; the response variable, network stability, and all three network-level predictor variables were mean-centered and standardized, so that the estimates would be comparable with the observed data (as summarized in the rightmost column of this table). The sample size for each analysis was 3,000 simulated networks. The first simulation models a scenario of reciprocity and behavioral heterogeneity using three simple rules: (i) reciprocal partner choice, (ii) a preference for previous social partners, and (iii) among-individual differences in the capacity for social behavior. The null model removed all three of these conditions. For comparison, we also present a second null scenario whereby stability was calculated after randomly rewiring the network. Finally, we considered additional scenarios based on (i), (ii), or (iii) alone. Notably, only the full simulation model that combined reciprocity and behavioral heterogeneity (i + ii + iii) was able to recapitulate the effects of network weight and density on stability. The individual-based simulations also demonstrated that network size has a negative effect on stability, with larger networks being less stable, except in the case of random rewiring where network size has no influence on stability.

| Simulation model | Predictor | Estimate (std. coef.) | Lower 95% confidence interval | Upper 95% confidence interval | t | p-value | Sign of the estimate | Within 95% CI of estimate from obs. data? |
|---|---|---|---|---|---|---|---|---|
| i + ii + iii | Network size | –0.40 | –0.41 | –0.38 | –54.05 | < 0.0001 | – | Y |
| | Network weight | 0.29 | 0.28 | 0.30 | 39.43 | < 0.0001 | + | Y |
| | Network density | –0.77 | –0.78 | –0.75 | –104.42 | < 0.0001 | – | Y |
| Null model | Network size | –0.28 | –0.29 | –0.27 | –32.15 | < 0.0001 | – | Y |
| | Network weight | 0.007 | –0.009 | 0.02 | 0.87 | 0.38 | n.s. | n |
| | Network density | 0.84 | 0.82 | 0.86 | 97.92 | < 0.0001 | + | n |
| Random rewiring | Network size | –0.003 | –0.01 | 0.007 | –0.54 | 0.59 | n.s. | Y |
| | Network weight | 0.001 | –0.009 | 0.01 | 1.66 | 0.87 | n.s. | n |
| | Network density | 0.96 | 0.95 | 0.97 | 191.99 | < 0.0001 | + | n |
| i only | Network size | –0.32 | –0.36 | –0.29 | –18.58 | < 0.0001 | – | Y |
| | Network weight | 0.009 | –0.03 | 0.04 | 0.51 | 0.61 | n.s. | n |
| | Network density | 0.01 | –0.02 | 0.04 | 0.63 | 0.53 | n.s. | n |
| ii only | Network size | –0.34 | –0.37 | –0.31 | –24.51 | < 0.0001 | – | Y |
| | Network weight | –0.002 | –0.03 | 0.03 | –0.12 | 0.90 | n.s. | n |
| | Network density | –0.55 | –0.58 | –0.52 | –39.65 | < 0.0001 | – | Y |
| iii only | Network size | –0.065 | –0.07 | –0.06 | –17.73 | < 0.0001 | – | Y |
| | Network weight | 0.06 | 0.05 | 0.06 | 15.50 | < 0.0001 | + | n |
| | Network density | 0.98 | 0.97 | 0.98 | 264.27 | < 0.0001 | + | n |
| Three time steps | Network size | –0.39 | –0.40 | –0.37 | –47.24 | < 0.0001 | – | Y |
| | Network weight | 0.28 | 0.26 | 0.30 | 34.17 | < 0.0001 | + | Y |
| | Network density | –0.76 | –0.77 | –0.74 | –92.67 | < 0.0001 | – | Y |
| Ten time steps | Network size | –0.42 | –0.43 | –0.40 | –63.22 | < 0.0001 | – | Y |
| | Network weight | 0.30 | 0.28 | 0.31 | 44.96 | < 0.0001 | + | Y |
| | Network density | –0.78 | –0.80 | –0.77 | –118.58 | < 0.0001 | – | Y |

**Table S4. Analysis of long-term partnership persistence.**

Posterior estimates from an analysis of the probability of partnership persistence from one year to the next (n = 669 partnerships involving 91 individuals). The model is a Bayesian multiple-membership analysis with a binary response variable (whether each social partnership, or network edge, persisted the following year). Field season is a categorical variable with only two levels (2015-16 and 2016-17), because persistence beyond 2017-18 is not known. The model posteriors indicate a substantial drop in edge persistence after the second season (2016-17), as compared to the first (2015-16). Status refers to status of interacting dyads (categorical with three levels: floater + floater, floater + territorial, and territorial + territorial). Status composition and edge betweenness were not associated with partnership persistence independent of the other predictors. Instead, bonds were more likely to persist if the two individuals had greater spatial overlap, if they interacted more frequently (edge weight), and if they had a lower edge connectivity within the social network. Sampling effort was quantified as the number of recording hours at the lek where the partnership primarily occurred, both in its initial year (“yr1”), and the following year when persistence was assessed (“yr2”). All continuous predictors were mean-centered and standardized to allow relative effect sizes to be compared across the table. The Bayesian $R^2$ for the fitted model is 0.36, indicating that over a third of the total variance in edge survival can be explained by this analysis.

| Fixed effect | Posterior estimate (std. coef.) | Standard error | Lower 95% credible interval | Upper 95% credible interval | Effective sample size |
|---|---|---|---|---|---|
| Field season | | | | | |
| 2016-17 | –1.34 | 0.53 | –2.37 | –0.27 | 3,050 |
| Status | | | | | |
| Floa. + Terr. | 0.43 | 0.38 | –0.29 | 1.19 | 3,344 |
| Terr. + Terr. | 0.11 | 0.52 | –0.92 | 1.14 | 2,944 |
| Sampling effort yr1 | 0.00 | 0.01 | –0.01 | 0.01 | 2,799 |
| Sampling effort yr2 | –0.00 | 0.01 | –0.01 | 0.01 | 4,665 |
| Spatial overlap | 0.33 | 0.11 | 0.13 | 0.54 | 3,308 |
| Edge weight | 0.80 | 0.12 | 0.58 | 1.04 | 3,910 |
| Edge betweenness | 0.03 | 0.11 | –0.19 | 0.24 | 5,232 |
| Edge connectivity | –1.11 | 0.39 | –1.88 | –0.36 | 5,091 |

**Table S5. Model selection for individual behaviors that predict long-term partnership persistence.**

Candidate models to evaluate the behavioral phenotypes that predict social fidelity (n = 91 males). The response variable was the posterior median for an individual's effect on annual partnership persistence. A male with a high value of this trait is one who promotes long-term stable partnerships, whereas a male assigned a low value of this trait does not. We fit candidate linear models that were ranked by AIC. The evidence ratio compares the support for model i to the best-fit model, and is calculated using the Akaike weights as $w_1/w_i$. The results indicate that two phenotypes, a male's average daily frequency of social interactions (strength) and his average daily number of social partners (degree), were significant but non-independent predictors of the stability of his partnerships in the subsequent year. Models with either of these two correlated behaviors received similar support, with positive effects of strength or degree that explained 11% of the variance in persistence as an individual trait.

| Rank, i | Predictors | $AIC_i$ | $\Delta AIC_i$ | $w_i$ (AIC) | Evidence ratio (best-fit vs. model i) | $R^2$ |
|---|---|---|---|---|---|---|
| 1 | Strength (daily avg.) + Status + Sampling eff | 173.49 | 0 | 0.53 | – | 0.11 |
| 2 | Degree (daily avg.) + Status + Sampling eff | 174.35 | 0.86 | 0.34 | 1.5 | 0.11 |
| 3 | Effort (daily avg.) + Status + Sampling eff | 177.71 | 4.22 | 0.06 | 8.8 | 0.07 |
| 4 | Effort + Strength + Degree + Importance + Status + Sampling eff | 178.68 | 5.19 | 0.04 | 13.3 | 0.12 |
| 5 | Status + Sampling eff | 180.49 | 7.00 | 0.02 | 33.1 | 0.02 |
| 6 | Importance (daily avg.) + Status + Sampling eff | 182.49 | 9.00 | 0.01 | 88.3 | 0.02 |

**Table S6. Analysis of the trade-off between sociality and partnership maintenance.** We used a repeated measures analysis to partition the variance in social stability within- and among-individuals. The response variable, partnership maintenance, is defined as the proportion of social ties that a given male maintained from one recording session to the next (n = 565 observations of 152 individuals). The binomial mixed-effects regression model included a random effect of male identity to account for repeated measures, as well as fixed effects to account for among-season variation (categorical with three levels: 2015-16, 2016-17, 2017-18), status class (categorical with two levels, floater or territorial), and sampling effort (continuous, based on the number of recording hours per territory in the recording session at the male's primary lek). The analysis also partitioned the among- and within-individual variance by considering average and relative degree, respectively, as the two fixed effects of interest. Both of these continuous predictors were mean-centered and standardized to allow their effect sizes to be compared. Among individuals, the more social males (i.e., those with more partners) exhibited significantly greater partnership maintenance, consistent with the analysis in Table S5. However, within individuals, a male's relative degree was negatively related to his subsequent partnership maintenance. This suggests that each male has constraint on the number of social ties that he can maintain at any one time, a trade-off that is not apparent at the among-individual level owing to among-individual heterogeneity.

| Fixed effect | Estimate (std. coef.) | Lower 95% confidence interval | Upper 95% confidence interval | z | p-value |
|---|---|---|---|---|---|
| Field season | | | | | |
| 2016-17 | 0.13 | –0.06 | 0.32 | 1.31 | 0.19 |
| 2017-18 | 0.19 | –0.02 | 0.41 | 1.75 | 0.08 |
| Status | | | | | |
| Territorial (vs. Floater) | 0.42 | 0.24 | 0.60 | 4.61 | < 0.0001 |
| Sampling effort | 0.15 | 0.07 | 0.22 | 3.87 | 0.0001 |
| Average session degree | 0.13 | 0.02 | 0.24 | 2.39 | 0.02 |
| Relative session degree | –0.35 | –0.45 | –0.25 | –6.89 | < 0.0001 |

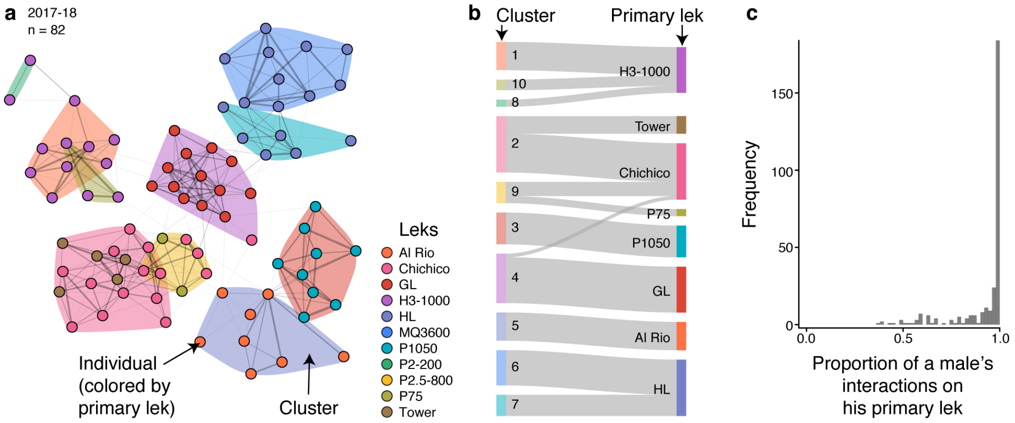


**Figure S1. Social structure corresponds to lek structure.** We performed a clustering analysis to verify that the lek units captured the natural social structure of the manakin population. The first step was to find the natural community structure of each annual social network (i.e., defined as the aggregation of all recording sessions and all leks in a given field season). To do this, we used the random walk clustering algorithm in the igraph package. The three annual social networks were highly modular (modularity coefficients of 0.70, 0.73, and 0.83, respectively), demonstrating strong clustering of interactions into communities. **a**, Example annual social network from 2017-18. Each node represents an individual male and is colored according to his primary lek, defined as the site where he engaged in the most cooperative interactions. The overlaid color polygons indicate cluster membership. The layout of the network was determined using the force-directed LGL algorithm, with edges weighted by the interaction frequencies. Next, we compared annual cluster membership to each male's primary lek designation, defined as the lek where the male engaged in the greatest number of social interactions that year. The probability that a male's primary lek matched the majority of his cluster was 90% (266/296 male-years). **b**, Example Sankey diagram to compare the clustering algorithm with primary lek usage. The relatively low proportion of crossings demonstrates strong correspondence. **c**, Histogram showing how males typically limited nearly all of their interactions to a single lek. This spatial patterning of interactions creates structure (modularity) in the network, resulting in the strong correspondence between lek structure and the clustering algorithm. Note that the same color schemes are used throughout **a–b**. Finally, the assortativity coefficient provides another way to compare lek and cluster structure in the social networks. The three annual networks were highly assorted by primary lek (coefficients 0.51, 0.51, and 0.69) to a level that was statistically indistinguishable from the assortment based on cluster membership (coefficients 0.47, 0.52, and 0.69, respectively). Based on this strong correspondence between lek structure and the structure revealed by the clustering algorithm, we conclude that our sampling design was well-matched to the inherent social structure of the population.

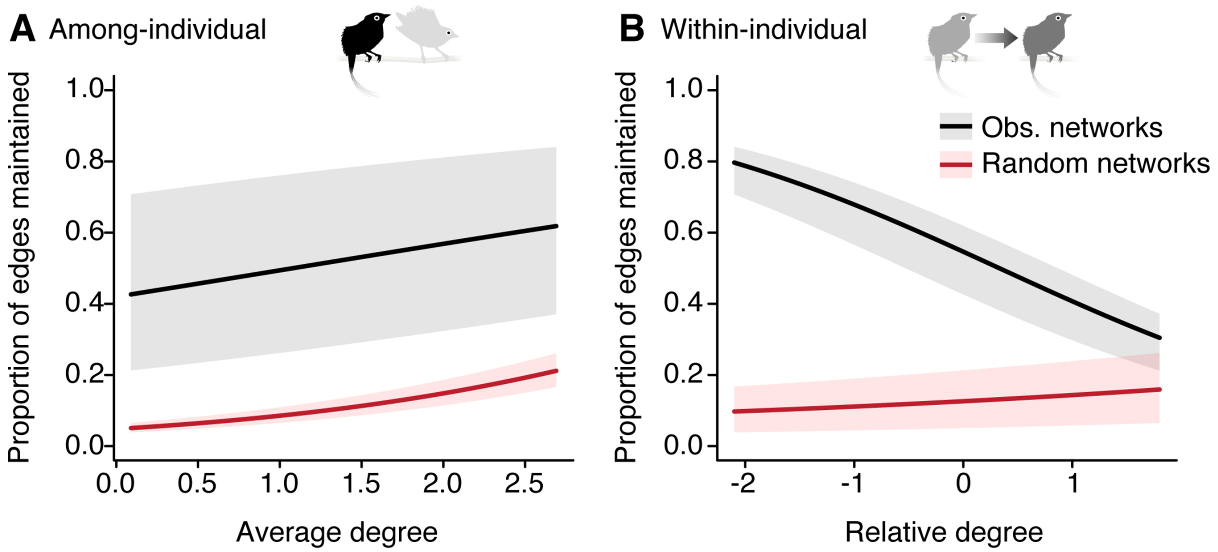


**Figure S2. Null expectation for the trade-off analysis in Table S6.** The results in shown in gray illustrate the trade-off analysis of the manakin data from Fig. 4B-C, as compared to a scenario shown in red where the social network dynamics are instead determined by random rewiring. In (A), we see that even though a weak positive association between a male's average degree and his edge maintenance is expected to occur by chance, it is notable that edges are in general much more likely to be maintained in the real networks as opposed to the random scenario. In (B), we see that the strong negative effect of relative degree observed in the manakin networks is not observed in the random scenario. Instead, under random rewiring, a weak positive effect is expected. The lines-of-best-fit show the predictions from repeated-measures analyses when all other fixed effects were set to their mean values. Each shaded region was determined by setting the other degree parameter to its minimum and maximum value, respectively. For the null expectation in red, results were obtained by averaging across the analyses of 1,000 randomly permuted datasets.

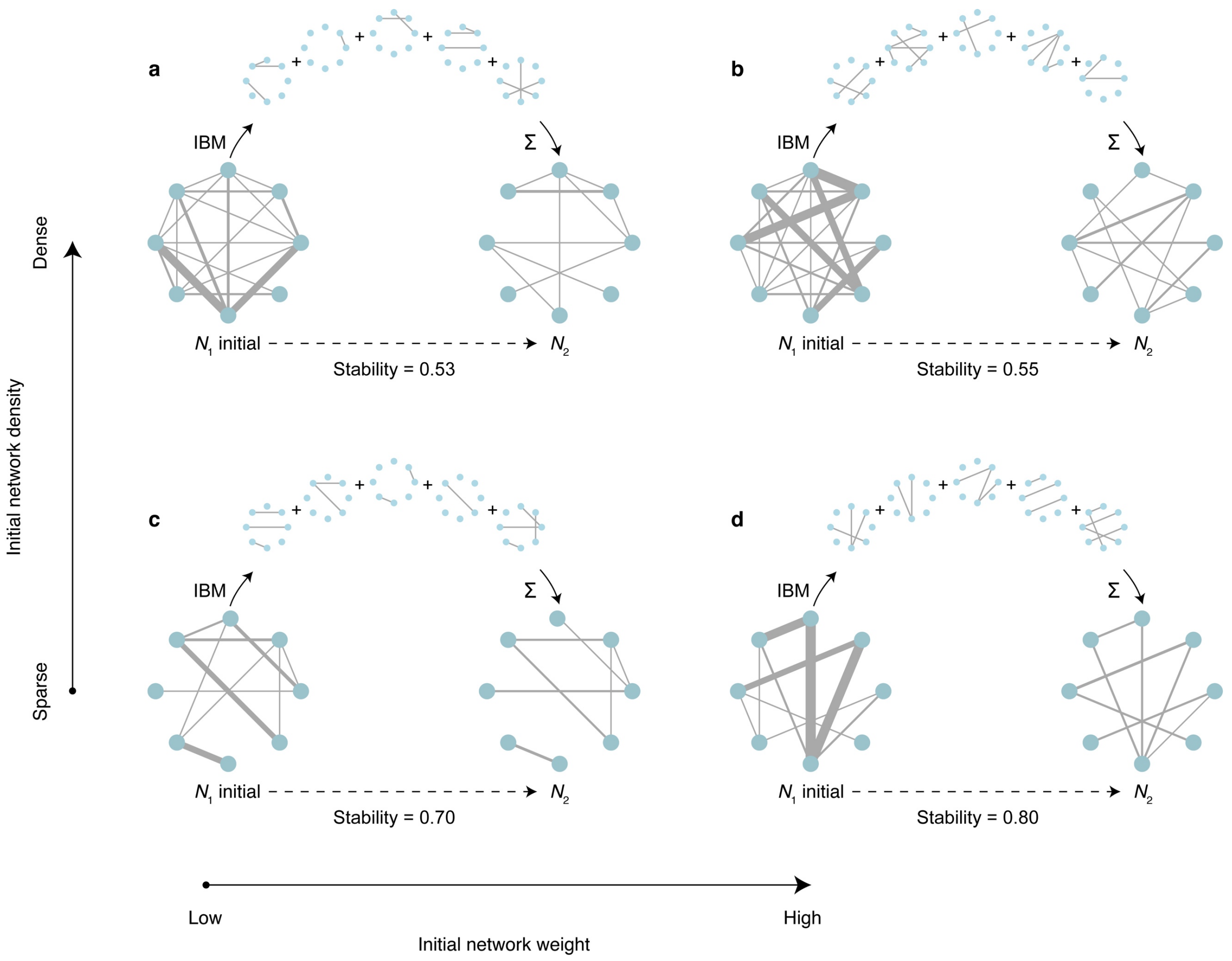


**Figure S3. Procedure for the individual-based simulation models. a–d**, Four examples illustrate the general approach and the data generated. The initial social network ($N_1$) represents the starting point used to determine the probabilities of subsequent social interactions. Each individual-based model (IBM) was run for five time-steps, akin to measuring interactions across multiple days. During this time, new partnerships were formed, as shown. The subsequent network ($N_2$) represents the sum of new interactions that occurred during the IBM time-steps. The stability score is calculated by comparing the topology of $N_2$ with that of $N_1$. Note that within each example **a–d**, the layout of particular nodes is kept constant throughout. Edge thicknesses are scaled to the log-transformed interaction frequency in a consistent manner throughout the figure. Note that the $N_2$ networks have a lower network weight as compared to $N_1$, because the time interval for the simulation was relatively short (albeit sufficient to develop a wide range of network stability values). The four examples **a–d** were chosen because they vary in the initial network weight (increasing left to right) and density (increasing bottom to top). The network size of eight was used in this figure for visual clarity in the illustration; however, in the actual simulations, network sizes ranged from 11-20 individuals.
